# Dynamics of endogenous and water cortisol release in Asian Seabass *Lates calcarifer* after acute stress in a farm scale recirculating aquaculture system

**DOI:** 10.1101/2023.09.20.558587

**Authors:** Marie Ruoyun Tan, Khin Moh Moh Aung, Nur Asinah binte Mohamed Salleh, Jolin Yean Ai Tan, Kai Xin Chua, Gaynah Javier Doblado, Kai Lin Chua, Valarie Tham, Jovian Jing Lin, Vindhya Chaganty, Dinah Mardhiana Yusoff, Shubha Vij, Xiaodi Su, Laura Sutarlie, Caroline Lei Wee

## Abstract

Stress in farmed fish is associated with poor feeding, slow growth, disease, and mortality. Therefore, it is essential to closely monitor the stress levels in fish to optimize farming practices which could then enhance productivity and welfare in aquaculture operations. Cortisol, a stress hormone that can be found in the blood, is a reliable biomarker for evaluating fish stress. As blood sampling is highly invasive, alternative cortisol sampling methods such as fin, mucus, and the surrounding water which contains released cortisol, have been proposed as less invasive or non-invasive sampling methods. However, a comprehensive understanding of their temporal dynamics and associations with plasma cortisol levels is still lacking. In this study, we subjected *Lates calcarifer*, Asian sea bass within a farm-scale (3,000 L tank, 9,000 L system) high-flow rate (8,000 L/hour) Recirculating Aquaculture System (RAS) to an acute handling stress challenge specifically involving chasing and air exposure, and quantified cortisol dynamics both within different biological samples including blood, fin, and mucus and in tank water from multiple sampling points. We showed that handling stress induced an expected increase in plasma and mucosal cortisol, peaking at 1 hour and 24-48 hours, respectively, and that plasma and mucus cortisol were moderately correlated, especially during the stress period. Fin cortisol did not show consistent dynamics. Water cortisol similarly rose, but peaked within 40 minutes from the start of the stressor, in a pattern that was dependent on the site of sampling within the RAS system, likely due to RAS circulation dynamics. Our study is the first to examine the impact of stress on cortisol accumulation and release in Asian Sea bass in a farm-scale RAS, thus complementing existing research on the efficacy of fin, mucus, and water cortisol as stress indicators that could help optimize aquaculture productivity and welfare.

## Introduction

Fish stress has negative effects on growth, survival, and meat quality, and is both a cause and consequence of disease (Snieszko, 1974; Bly, Quiniou and Clem, 1997; Daskalova, 2019). Pathogenic infections (Ellis *et al*., 2007; Triki *et al*., 2016) and environmental stressors such as poor water quality (Lupica and Turner, 2010; Mota *et al*., 2017a; Zarantoniello *et al*., 2021), handling (Scott, Pinillos and Ellis, 2001; Ellis *et al*., 2004), noise (Mickle and Higgs, 2018), and overcrowding (Pavlidis *et al*., 2013; Odhiambo *et al*., 2020) commonly induce the stress response in fish, which is marked by the secretion of the stress hormone, cortisol. As such, cortisol has been utilized as a biological indicator of fish stress levels in both laboratory and aquaculture settings (Martínez-Porchas, Martínez-Córdova and Ramos-Enriquez, 2009; Sadoul and Geffroy, 2019; Tanaka *et al*., 2023). Unfortunately, conventional methods of obtaining cortisol from fish blood, or whole-body samples are invasive and themselves stress-inducing (Scott, Pinillos and Ellis, 2001; Sadoul and Geffroy, 2019).

Measurements of cortisol concentrations from mucus and fin have been used as less-invasive methods for stress monitoring (Simontacchi *et al*., 2008; Bertotto *et al*., 2009; De Mercado *et al*., 2018; Ghassemi Nejad *et al*., 2019). Simontacchi et al (2008) evaluated plasma cortisol levels in European sea bass subjected to different pre-slaughter conditions. Cortisol levels in plasma were found to correlate with those detected in mucus, even though cortisol levels in mucus were substantially (up to 20 times) lower than in plasma. Bertotto et al (2010) investigated transportation stress in European sea bass and other fishes and similarly observed a good correlation between cortisol levels in plasma, mucus and fin samples.

Cortisol within the animal’s bloodstream is also released into the surrounding water through passive diffusion (Sadoul and Geffroy, 2019), and water cortisol levels were previously shown to correlate with plasma cortisol levels (Scott, Pinillos and Ellis, 2001; Fanouraki *et al*., 2008). Hence, regular measurement of cortisol in the water has been proposed as a non-invasive means of detecting elevated stress levels in farmed fish (Scott, Pinillos and Ellis, 2001; Sadoul and Geffroy, 2019). This method might be particularly useful in land-based farms that use Recirculating Aquaculture Systems (RAS) technologies for high-intensity farming (Mota *et al*., 2017a).

A handful of previous studies have looked into the potential of water cortisol as a non-invasive stress marker. Fanouraki et al (2008) subjected European Seabass (*Dicentrarchus labrax*) to 5 minutes of chasing and a 1-1.5 minutes air exposure, within a 2 m^3^ flow-through tank, at a water flow rate of 60 L/hour (Fanouraki *et al*., 2008). This resulted in increased plasma cortisol concentrations (2,184 nM or 791,612 ng/L) that peaked at 1-hour post stress. Water cortisol concentrations peaked at 4-hours post-stress reaching 7.2 ± 0.20 ng/L. In the same study, a confinement challenge was also conducted with two stocking densities (20 or 50 kg/m^3^) within 12 L buckets with supplied O_2_ and a high water flow rate. The higher stocking density tank was observed to have had a 2-3 fold higher water cortisol concentration throughout the 24-hour experiment. In a different study, Scott et al (2001) showed that 90-second air exposure of rainbow trout (*Oncorhynchus mykiss*) induced stress as measured from water cortisol (Scott, Pinillos and Ellis, 2001). The study was conducted in a 150 L flow-through tank, with a stocking density of 30 kg/m^3^ at a water flow rate of 120 L/hour. A progressive increase in water cortisol concentrations was observed after single and repeated stress, peaking at 2 hours (25 ng/L) and 5 hours (100 ng/L) respectively, and gradually declining thereafter. The above studies used small volume, flow-through setups not comparable to farm-scale conditions.

More recently, water cortisol experiments have been conducted in RAS systems, albeit in relatively small setups. Mota et al (2017) examined the effect of RAS flow rate on water cortisol concentrations prior to and following an acute stressor (Mota *et al*., 2017b). Nile Tilapia (*Oreochromis niloticus*) were raised at a stocking density of 67 kg/m^3^ in 72 L tanks. After two weeks at a flow rate of 675 L/kg feed/day, water exchange rates were adjusted to 150 L/kg feed/day (LowRAS) or 1,500 L/kg feed/day (HighRAS) for 4 weeks, where LowRAS expectedly led to an increase in water cortisol concentrations over 4 weeks. Cortisol concentration peaked at 2 hours following an acute stressor of 60-second air exposure, increasing 30% in LowRAS from 5 ng/L to 7 ng/L, unlike the HighRAS condition, where a significant change was not observed. Another study by the same authors investigated the water exchange rate and pH on circulating cortisol in rainbow trout (*Oncorhynchus mykiss*) and the accumulation of cortisol in a RAS tank over a 70-day experimental period, with a starting stocking density of around 10 kg/m^3^. On day 35, plasma cortisol was significantly elevated at low pH (5.8) with a mean concentration of 24.4 +/- 9.5 ng/mL, and elevated water cortisol (2.5 ng/L per kg) was also observed at low water exchange rates (480 L/kg feed/day). These studies suggest that the flow rate of a RAS tank may significantly impact the ability to detect water cortisol, and a lower rate is essential for ensuring the utility of water cortisol as an on-farm stress diagnostic tool.

The Asian sea bass (*Lates calcarifer*), also known as Barramundi, is a commercially important fish species widely distributed in the Indo-Pacific region (Islam *et al*., 2023). It holds a prominent position among the marine species farmed in Singapore because of its consumers’ high demand, sustainability, economic benefits, palatable taste, and nutritional value (Glencross, 2006; Taylor, 2022). In aquaculture farming, Asian sea bass have a low feed conversion ratio, which means it requires less feed to produce one kilogram of fish as compared to other aquaculture species (Katersky and Carter, 2005). Additionally, it is commonly raised in RAS, which minimizes the environmental impact and reduces the risk of diseases and parasites, making it a sustainable fish to farm, especially in countries with limited land and water supplies. Overall, this high-value food fish generates revenue for fish farmers, as well as for local restaurants and markets (Keat, 2021). Similar to other farmed fish, Asian sea bass are often subjected to stress during various aquaculture practices, such as transportation, handling, and stocking. Therefore, stress detection in Asian sea bass is of significant interest to both the fisheries and aquaculture industries. However, few studies have been carried out on stress in Asian sea bass (Ardiansyah and Fotedar, 2016; Hong *et al*., 2021), and no quantification of cortisol accumulated or released into multiple tissue types or water has been reported.

Hence, in this study, we sought to characterize cortisol dynamics of Asian sea bass in response to stressors, particularly whether fin and mucus tissues may allow for accurate and less-invasive proxies of stress in this species. Furthermore, we investigated if water cortisol levels within a larger, farm-like, high-flow RAS setting (9,000 L system, 8,000 L/hour flow rate) would be an accurate measure of fish stress within the system, and compared water cortisol concentrations across different sampling points in the RAS system.

## Materials and Methods

### Animal Husbandry and Experimental Setup

Asian sea bass (*Lates calcarifer*) were reared in a RAS system at the Aquaria of Republic Polytechnic, Singapore, and all experiments were approved by the Republic Polytechnic’s Institutional Animal Care and Use Committee (IACUC Protocol #2022/RP/00001). Chasing experiments were carried out in a 9,000 L saltwater (30 ppt) system, which consisted of 2 circular blue fiberglass tanks (each 3,000 L) containing the fish and 1 rectangular tank (3,000 L) as the sump (Figure 1A-B). Water was circulated at 8,000 L/hour.

**Figure 1:**
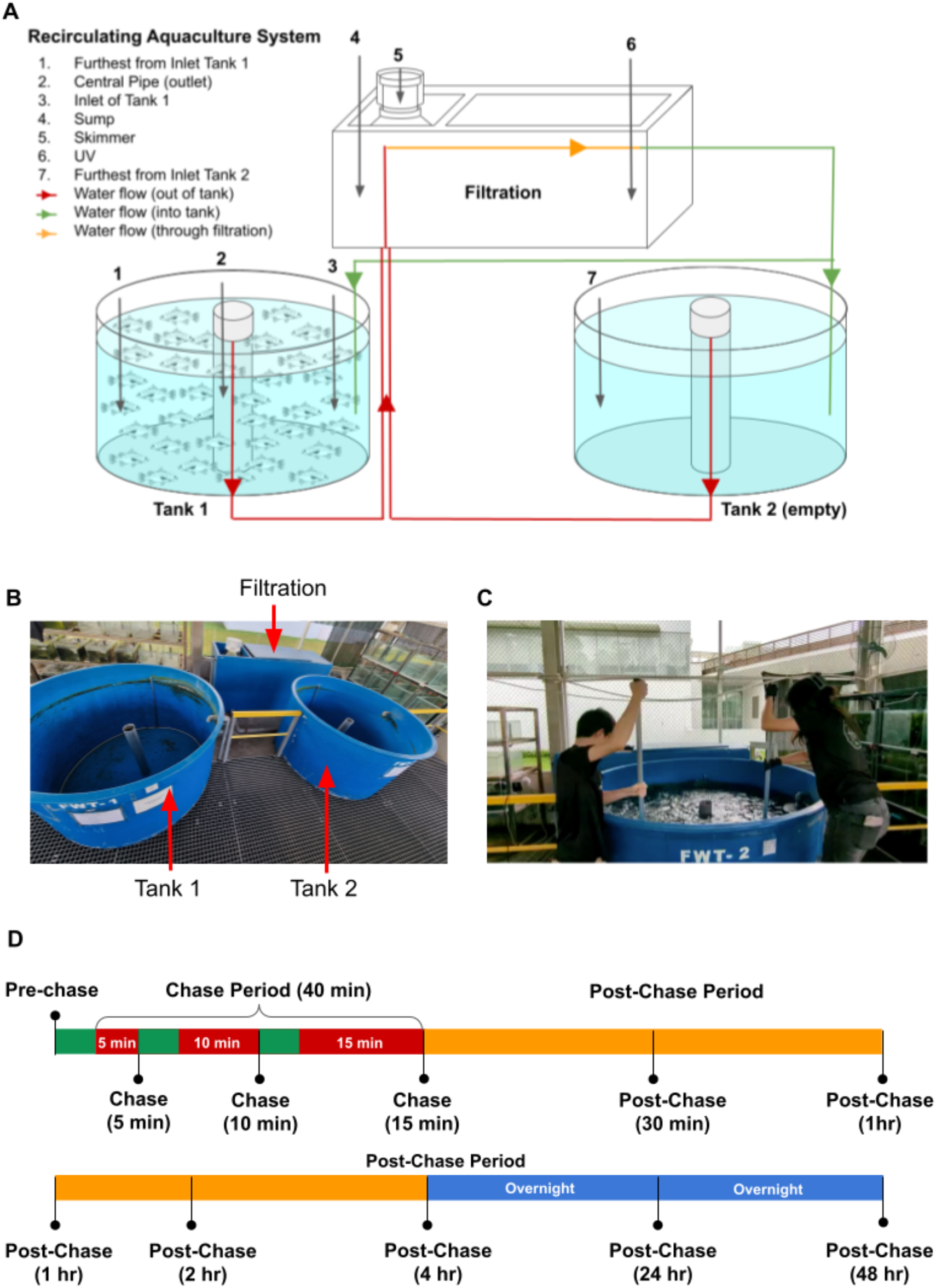
Schematic of experimental setup and design. (A) Design of the Recirculating Aquaculture System (RAS) used in our experiments, including water sample collection points. (B) Image of the RAS (C) Image showing how chasing was performed in Tank 1 using large nets in a C-shape manner. Air Exposure for as many fish as possible was performed for 10 s at the end of each chasing period (5 min, 10 min, 15 min) before 5 fish were sampled per time point. (D) Experimental timeline from Pre-Chase to Post-Chase (48 hr). Black lines correspond to sampling time points. Red boxes represent progressively longer stress periods separated by 5-minute intervals (green boxes). Orange boxes represent the post-stress period on the same day, following which sampling was also done after overnight rest periods (blue boxes).

### Experimental Fish

Asian sea bass were obtained from two local commercial fish farms. Upon arrival, the juvenile Asian sea bass (∼30 g) were subjected to a two week quarantine period to facilitate adaptation and address any diseases that the fish might have brought. Each batch of fish was placed directly and quarantined in a single fiberglass RAS tank “Tank 1” (Figure 1A-B). Details of each batch of fish used in the experiments can be found in Table 1.

**Table 1:** Summary of Asian sea bass used in experiments.

| Experiment Details |  |  | At start of experiment |  |  |  |
| --- | --- | --- | --- | --- | --- | --- |
| Biological Replicates | Start Date | Fish Supplier | Number of fish | Average Weight (g) | Stocking density (system, kg/m<sup>3</sup>) | Stocking density (tank, kg/m<sup>3</sup>) |
| Exp 1 | 7 Sep - 9 Sep 2022 | Local Farm A | 134 | 140.67 | 2.094 | 6.283 |
| Exp 2 | 7 Dec - 9 Dec 2022 | Local Farm B | 446 | 92.36 | 4.575 | 13.731 |
| Exp 3 | 1 Mar - 3 Mar 2023 | Local Farm B (same batch as Exp 2) | 432 | 165.82 | 7.959 | 23.878 |

During experimental periods, fish were fed three times a day (9:30 am, 12:30 pm, and 4:30 pm) at a rate of about 10% body weight (BW) per day using a commercial fish diet (5 mm sinking pellets; 46% crude protein, 10% crude fat, 5% crude fiber, 16% crude ash, 4.5% phosphorus, and 2.2% lactic acid). The amount of feed was also controlled accordingly due to the recirculating aquaculture system filter carrying capacity and fish stocking densities.

Water quality parameters were measured daily by the YSI Pro Quattro probe (YSI, USA) and the SpinTouch FX (Lamotte, USA). The 9,000 L RAS system that was used for chasing stress experiments was circulating aerated saltwater (dissolved oxygen, 5.74 ± 4.03 mg/L; water temperature, 27.58 ± 3.8 °C; salinity maintained at 30 ppt). Ammonia levels were controlled to be below 1 ppm, nitrite concentrations were maintained below 1 ppm, nitrate levels were kept below 100 ppm and pH was maintained below 8.0.

### Handling Stress experiments

Fish were starved and not fed during the evenings the day before experiments commenced. “Pre-chase” samples were collected before any chasing stresses were induced on the fish. Fish were subjected to a period of chasing with progressively longer chasing durations: 5 minutes (“Chase (5 min)”), 10 minutes (“Chase (10 min)”) and 15 minutes (“Chase (15 min)”), with a 5-minute “rest” interval between each of the 3 chases. Fish were chased by two personnel using large nets (Figure 1C), each in a repetitive “C” shape manner hence covering the entire breadth and depth of the tank. At the end of each of the respective chases, a net full of fish was lifted in one stroke and air exposed for 10 seconds. Following this, 5 fish from this net were collected for sampling.

Post-chase samples were collected at 30 minutes (“Post-chase (30 min)”), 1 hour (“Post-chase (1 hr)”), 2 hours (“Post-chase (2 hr)”), 4 hours (“Post-chase (4 hr)”), 24 hours (“Post-chase (24 hr)”), and 48 hours (“Post-chase (48 hr)”), from the end of the chasing period.

Hence, there were a total of 11 sampling time points for chasing experiments; where 5 random fish were sampled at each sampling time point (Figure 1D). At each sampling point, the following samples and data were collected: water from various parts of the tank (see “Water cortisol collection” section), fish body weight (total weight of the 5 fish), plasma, fin, and mucus (See “Biological sample collection” section).

### Biological sample collection

After being netted, the fish were anesthetized in a bucket using Tricaine methanesulfonate MS-222 (70 mg/L) and sodium bicarbonate (0.14 g/L) dissolved in fresh saltwater from the reservoir. While still in the bucket, the fish were collectively weighed. The anesthesia took effect in approximately 5 minutes, after which the fish were subjected to plasma, fin, and mucus collection, as described in the following sections. On average, the entire process of collecting plasma, fin, and mucus samples required approximately 5 minutes per fish.

### Plasma collection

Blood was extracted from the caudal artery/vein by inserting a 21G sterile needle at a 45° angle until it reached the spine. Upon contact with the spine, the needle was withdrawn slightly, and blood was extracted into a sterile 1.0 mL syringe. Approximately 0.5 mL of the collected blood was subsequently injected into a heparin tube and inverted 5 times to thoroughly mix the blood within the heparin tube and prevent coagulation. The blood in the heparin tube was then centrifuged at a 45° angle, at 3,000g for 10 minutes to facilitate the separation of blood plasma. The plasma was carefully pipetted into a cryovial, which was then stored in dry ice before being transferred to a -20 °C freezer.

### Mucus collection

The fish were swabbed on their left flank, along the lateral line. This process was repeated 4 times on each side of the swab. The swab was then placed into a 2 mL tube, where 1 mL of phosphate buffer solution (PBS) was added to the sample swab. The samples were then stored in dry ice before being transferred to a -20 °C freezer.

### Fin collection

A fin clip was collected by cutting about 1.5 cm of the top part of the caudal fin. Tweezers were then used to pick up the fins to be placed into 2 mL labeled Eppendorf tubes. The Eppendorf tubes with the fins were then filled with 1 mL of PBS and stored in dry ice before being transferred to a -20 °C freezer. After handling each fish, both tweezers and scissors were sanitized with 70% ethanol.

After sample collection, the fish were first placed in recovery saltwater buckets with no tricaine to recover from the anesthesia. This measure was implemented to prevent the resampling of the same fish at each sampling time and to avoid any potentially stressful effects caused by the reintroduction of the sampled fish. In the first experiment where water cortisol was only measured from “Tank 1 - Furthest from inlet”, the sampled fish were placed into “Tank 2” post-recovery from anesthesia. The latter was the other blue fiberglass tank connected to the entire recirculating system. In subsequent experiments, given that we were also sampling water from other parts of the system including Tank 2, the sampled fish were placed into a separate holding tank not connected to the circulating system and only returned at the end of the day after the “Post-chase 4 h” time point (Experiment 2), or in the case of Experiment 3, these fish were culled. We note that there were minimal changes in the system stocking densities caused by these differences (Supplementary Table 1), and as reported in the “Results” section, these minor variations did not appear to affect the overall trend of biological or water cortisol changes.

### Water sample collection

In Experiment 1, tank water was only collected from a single sampling point “Tank 1 Furthest from Inlet”. In Experiments 2 and 3, tank water was additionally sampled from 6 other sampling points (Figure 1A). Besides 1 - “Furthest From Inlet Tank 1”, water was sampled from the 2 - “Central Pipe”, 3 - “Inlet Tank 1”, 4 - “Sump”, 5 - “Skimmer”, 6 - “UV” compartment and 7 - “Furthest From Inlet Tank 2”. Gloves were consistently worn during the collection of water cortisol samples to avoid potential contamination from human skin-derived cortisol. Water was collected using siphons, positioned at a height that was midway through the water column. These siphons were thoroughly rinsed with clean freshwater at each respective collection point. To ensure thorough flushing of the siphon at each sampling point, the tank water was allowed to flow through the siphon for at least 30 seconds before a 100 mL or 125 mL glass bottle was filled to the brim. These water samples were subsequently kept chilled in an ice box and later transferred to a 4°C fridge for storage. Water from the system that had been siphoned out but not collected in the glass bottles was returned into the system to maintain a relatively constant water volume within the system.

### Cortisol extraction from fin and mucus samples

Sample processing was performed according to protocols adapted from previous methods (Ghassemi Nejad *et al*., 2019) with some modification. Mucus samples stored in a -20 °C freezer, were thawed to room temperature before processing. The microcentrifuge tube containing the swab with mucus was vortexed for 30 seconds, after which, the mucus was scrubbed from the swab with the wall and rim of the tube repeatedly. Following this, the solution was filtered using a 0.22 µm Polyvinylidene fluoride (PVDF) syringe filter and stored at a 4 °C fridge for analysis within a 24-hour timeframe.

Fin samples in PBS stored in a -20 °C freezer were thawed to room temperature before processing. Each fin sample was washed twice with isopropanol and subsequently air-dried for 2-3 days. The dried fin sample was ground into powder by using a mortar and pestle, and the weight of the ground powder was recorded. To facilitate cortisol extraction, 1.5 mL of methanol was added to the ground powder and shaken over a 3-day period. After centrifugation at 9,500 rpm for 10 minutes, the resulting supernatant was collected and placed in an oven at 38 °C for methanol evaporation. The extracted cortisol was reconstituted in 400 μL of PBS, filtered through a 0.22 µm PVDF syringe filter, and stored at 4°C fridge for analysis within 24 hours or in a -20 °C freezer for long-term storage.

### Cortisol quantification from biological samples (ELISA)

The cortisol levels from the three biological substrates (plasma, fin and mucus) were quantified using an enzyme-linked immunosorbent assay (ELISA) kit (Fish Cortisol ELISA, CUSABIO, Houston, TX, USA). The ELISA is based on the competitive inhibition enzyme immunoassay principle, involving the interaction between pre-coated cortisol and cortisol within the samples, and has a cortisol detection range of 0.0023–10 ng/mL.

Plasma and aliquots of cortisol extracted from fins and mucus were subjected to analysis following the protocol provided by the manufacturer. Briefly, all reagents and samples were brought to room temperature prior to use. Plasma samples were diluted 100 times with the provided sample diluent before testing. Extracted cortisol from fin and mucus samples were tested as these were without further dilution. In each well, 50 μL of sample or standard was mixed with 50 μL of 1× antibody and incubated at 37 °C for 40 minutes. Each well was then washed three times with the wash buffer and 100 μL of Horseradish peroxidase conjugated secondary antibody (HRP conjugate) was added immediately. The plate was incubated in the dark at 37 °C for 30 minutes and subjected to five additional washes. Color development was initiated by adding 90 μL of 3,3’,5,5’-Tetramethylbenzidine (TMB) substrate to each well and incubating the plate at 37 °C for 20 minutes. The reaction was then stopped by adding 50 μL of stop solution. The plate was gently tapped to ensure thorough mixing. The optical density of each well was determined within 5 minutes using a microplate reader, the Infinite M200 Spectrophotometer (Tecan Trading AG, Switzerland) at a wavelength of 450 nm. The cortisol concentration in the sample is calculated based on the standard curve, prepared by plotting the optical density at 450 nm of cortisol standards (0 - 10 ng/mL) derived from serially diluted cortisol stock (10 ng/mL).

### Cortisol extraction and quantification from water samples (HPLC)

Cortisol from tank water samples was extracted through liquid-liquid extraction technique and analyzed by using high-performance liquid chromatography (HPLC) (Viljoen *et al*., 2019; Ney *et al*., 2021) with some modifications. For liquid-liquid extraction of cortisol from water samples, equal volumes of dichloromethane and water samples were mixed together and the mixture was left to stand for 10 minutes. The organic phase containing cortisol was decanted from the mixture, and evaporated using a rotary evaporator (IKA RV 10, Germany). The resulting precipitate was then reconstituted in a 50% methanol solution and filtered through a 0.22 µm PVDF filter.

The cortisol extracted from water samples was analyzed using HPLC (Shimadzu LC-2050C 3D, Japan) coupled to a photodiode array (PDA) detector set at 245 nm. The HPLC was performed by injecting 100 µL of the extracted water sample through a Shim-Pact GIST C18 normal-phase column (inner diameter 4.6mm, length 250 mm, particle size 5 µm). An isocratic flow of mobile phases (methanol and 10 mM ammonium formate in water in a 1:1 ratio) was performed for 30 minutes. Using the HPLC conditions mentioned above, cortisol was eluted with a retention time of 14.5 minutes. The HPLC is calibrated to cortisol standards ranging from 1 pM to 10 µM in 50% methanol solution. All water samples and standards were spiked with 500 µL of 1 µM cortisol internal standard. Cortisol peak area from the standards was measured and plotted against the cortisol standards concentration to build a standard curve. The cortisol concentration in the water sample is calculated based on the standard curve according to the peak area in each water sample. Each water sample was measured as duplicates.

### Data analysis and Statistics

Cortisol concentrations obtained from plasma, fin and mucus samples were calculated based on a standard curve run on each plate and expressed in ng/L. Adjustments were made for dilution factor (plasma), initial sample weight (100 mg of fin/1 mL of PBS) and mucus dissolved in 1 mL PBS. In Figure 3, the water cortisol concentration was not standardized to biomass. However, a standardization to system stocking density (kg/m^3^) was performed in Supplementary Figures 4-5.

**Figure 2:**
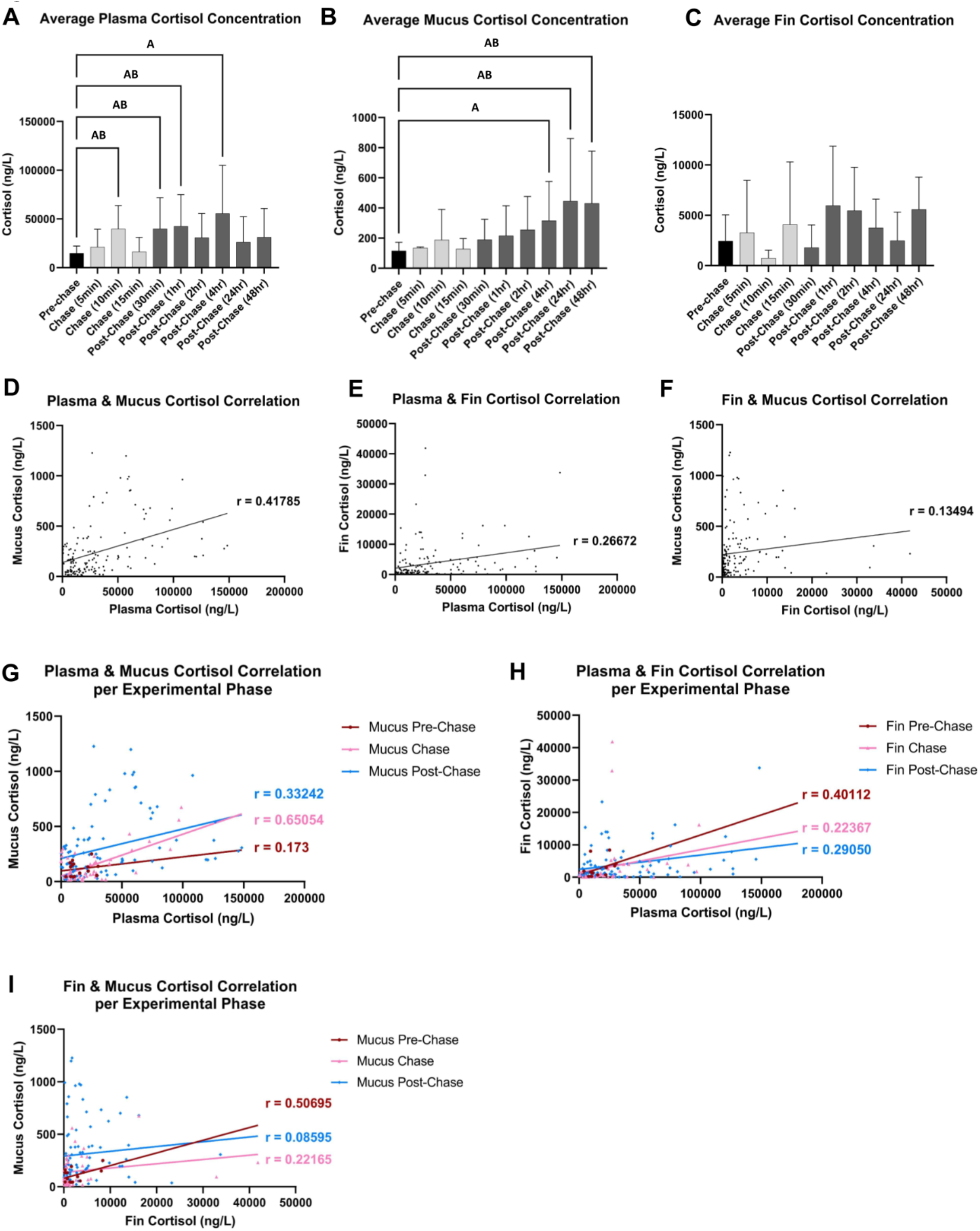
Changes in plasma, mucus, and fin cortisol levels during and after a period of chasing and air exposure stress. (A) The average plasma cortisol concentrations of Asian sea bass across sampling points (N = 5 fish per experiment, 15 fish per sampling point except for Chase (15min) where one outlier was excluded). A indicates statistical significance with two-way ANOVA only, whereas AB indicates statistical significance with both ANOVA and Bootstrapping analysis. P-values are reported in Table 2. (B) The average mucus cortisol concentrations of Asian sea bass across sampling points (N = 5 fish per experiment, 15 fish per sampling point). P-values in Table 2. (C) The average fin cortisol concentrations of Asian sea bass across sampling points (N = 5 fish per experiment, 15 fish per sampling point except for Pre-Chase and Post-Chase (24hrs) where one outlier was excluded each). P-values in Table 2. (D) Correlations between plasma and mucus cortisol concentrations released from individual fish across all time points (N = 148, R2 = 0.1746, p <0.0001) (E) Correlations between plasma and fin cortisol concentrations released from individual fish across all time points (N = 147, R2 = 0.07114, p = 0.0011) (F) Correlations between mucus and fin cortisol concentrations released from individual fish across all time points (N = 147, R2 = 0.01821, p = 0.1032) (G) Correlations between plasma and mucus cortisol concentrations released from individual fish per experimental phase (Pre-Chase: N = 15, R2 = 0.02993, p = 0.5375. Chase: N = 45, R2 = 0.4232, p <0.0001. Post-Chase: N= 89, R2 = 0.1105, p = 0.0015) (H) Correlations between plasma and fin cortisol concentrations released from individual fish per experimental phase (Pre-Chase: N = 14, R2 = 0.1609, p = 0.1552. Chase: N = 45, R2 = 0.05003, p = 0.1444. Post-Chase: N= 89, R2 = 0.08439, p = 0.0058) (I) Correlations between mucus and fin cortisol concentrations released from individual fish per experimental phase (Pre-Chase: N = 14, R2 = 0.2570, p = 0.0643. Chase: N = 45, R2 = 0.04913, p = 0.1434. Post-Chase: N= 88, R2 = 0.007388, p = 0.4259)

**Figure 3:**
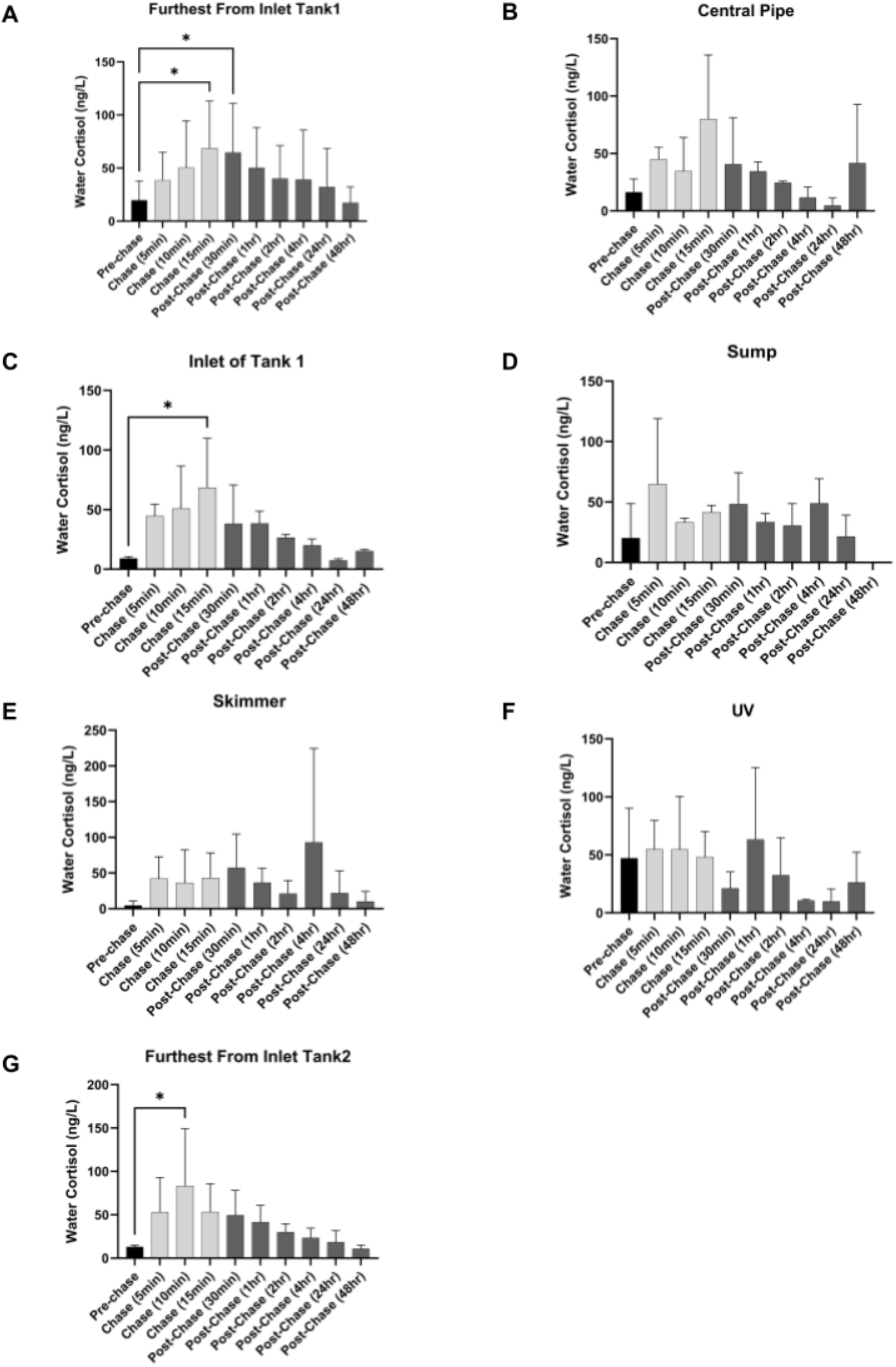
Change in water cortisol across different sampling points in the RAS system during and after a period of chasing and air exposure stress. (A) Mean water cortisol concentrations (ng/L) at Furthest From Inlet Tank 1 (N = 3 per time point) p = 0.0197 (Chase (15 min)), 0.0367 (Post-Chase (30min)) (B) Mean water cortisol concentrations (ng/L) at Central Pipe (N = 2 per time point) (C) Mean water cortisol concentrations (ng/L) at Inlet of Tank 1 (N = 2 per time point) p = 0.0287 (Chase (15 min)) (D) Mean water cortisol concentrations (ng/L) at Sump Tank (N = 2 per time point) (E) Mean water cortisol concentrations (ng/L) at Skimmer (N = 2 per time point) (F) Mean water cortisol concentrations (ng/L) at UV (N = 2 per time point) (G) Mean water cortisol concentrations (ng/L) at Furthest From Inlet Tank 2 (N = 2 per time point) p = 0.0395 (Chase (10min))

To compare cortisol concentrations across different time points, a 2-way ANOVA was applied, followed by the Dunnett test with multiple comparison correction. To ascertain the relationship between less-invasive measurements of mucus and fin cortisol and cortisol levels in blood plasma, we conducted simple linear regression analysis. Using the Grubbs’ test (Alpha = 0.0001) to identify outliers, we removed two fin cortisol (Experiment A, Pre-chase and Post-Chase (24hr)) and one plasma cortisol (Experiment A, Chase (15min)) measurement in our analysis.

Since the biological replicates are conducted on batches of fish with different characteristics and spanning six months, we also performed a boot-strapping based permutation test to establish the statistical robustness of our conclusions (MATLAB, USA). For 10,000 iterations, we first randomly sampled an experiment (1, 2, or 3), and from this experiment then randomly sampled (with replacement) 5 fish from each chase / post-chase time point. We then compared the mean cortisol concentration of this group with the pre-chase control, which was similarly randomly sampled. This allowed us to quantify the percentage of iterations in which the cortisol concentration would be higher in each post-chase time point compared to the pre-chase controls, providing a p-value for each time point. This permutation analysis was conducted for each tissue type. In this analysis, we did not exclude any outliers. Using this same analysis we also estimated the distribution of effect sizes (cohen’s d) that would be observed (Supplementary Figure 2). In the figures and tables, asterisks are used to indicate statistical significance levels as follows *p<0.05, **p<0.01, ***p<0.001, ****p<0.0001.

## Results

The acute handling stress experiment was repeated three times over a period of six months, as described in Table 1. Given its large scale, we were unable to simultaneously conduct multiple replicates, and instead conducted biological replicates sequentially over time. The origins, age, weight, and stocking densities of the fish inevitably varied, which led to varying absolute cortisol concentrations as quantified by ELISA analyses (Table 1, Supplementary Figure 1). However, handling stress and sampling protocols were kept consistent, and consistent behavioral changes were observed in each chasing experiment, where the fish exhibited rapid swimming, actively evaded the nets, and displayed signs of distress upon exposure to air. Instances of vomiting were observed, and foamy bubbles were observed on the surface of the tank, starting from around the “Chase (10 min)” period and increasing over time, which may reflect increased mucus secretion.

To quantify the effects of such handling stress on plasma, mucus, and fin cortisol across these diverse conditions, we employed two statistical methods, a 2-way ANOVA (across experiment and time point) as well as bootstrapping analysis. For the latter, we randomly sampled with replacement data from fish within each experiment, as well as the experiment (1, 2, or 3) from which the fish were sampled, for 10,000 iterations, to generate a bootstrapped distribution of effect sizes (Supplementary Figure 2) as well as a p-value for each time point (Table 2) demonstrating the probability of cortisol concentrations being higher at that time point relative to control.

**Table 2:** Summary of statistical analyses results using ANOVA and bootstrapping.

(dark gray = $p < 0.05$ in both tests, light gray = $p < 0.05$ in one test)
| Time points | Plasma |  | Mucus |  | Fin |  |
| --- | --- | --- | --- | --- | --- | --- |
|  | ANOVA | Bootstrap | ANOVA | Bootstrap | ANOVA | Bootstrap |
| Chase (5 min) | 0.9737 | 0.3374 | >0.9999 | 0.4147 | 0.9998 | 0.7656 |
| Chase (10 min) | 0.0183* | 0.0115* | 0.7128 | 0.6060 | 0.9673 | 0.5951 |
| Chase (15 min) | >0.9999 | 0.1726 | >0.9999 | 0.3560 | 0.9715 | 0.7236 |
| Post-chase (30 min) | 0.0172* | 0.0194* | 0.7181 | 0.2667 | >0.9999 | 0.8522 |
| Post-chase (1 hr) | 0.0066** | 0.0464* | 0.3574 | 0.3423 | 0.4202 | 0.3211 |
| Post-chase (2 hr) | 0.2616 | 0.1431 | 0.0803 | 0.1634 | 0.5990 | 0.3574 |
| Post-chase (4 hr) | <0.0001*** | 0.3089 | 0.0033** | 0.0718 | 0.9930 | 0.4320 |
| Post-chase (24 hr) | 0.6323 | 0.3656 | <0.0001*** | 0.0193* | >0.9999 | 0.8369 |
| Post-chase (48 hr) | 0.2317 | 0.1114 | <0.0001*** | 0.0000*** | 0.5471 | 0.3100 |

The results of both statistical analyses are tabulated in Table 2, and we only consider results that were significant across both statistical tests. Under this stringent criterion, we report a significant increase in plasma cortisol levels (Figure 2A), at the Chase (10 min), Post-chase (30 min and 1 hr) time points, with a complete return to baseline by 24 hours post-chase. In contrast, mucus cortisol showed a slower time course (Figure 2B), with a gradual increase of mucus cortisol over time, which only achieved significance at 24 and 48 hrs post-chase. Notably, mucus cortisol concentrations were about 100 fold lower than plasma or fin cortisol concentrations (Figure 2A-C). No significant values were obtained for fin cortisol (Figure 2C).

Next, we correlated cortisol levels across the biological samples (fin, mucus, and plasma) collected from each individual fish. Plasma cortisol was most strongly correlated with mucus cortisol (r = 0.42, p <0.0001, Figure 2D), whereas both fin and plasma (r = 0.27, p = 0.0011, Figure 2E), and fin and mucus cortisol (r = 0.14, p = 0.1032, Figure 2F), were only weakly correlated. Notably, the correlation between plasma and mucus cortisol was highest during the chasing period (r = 0.65, p <0.0001) as compared to the pre-chase (r = 0.33, p = 0.5375) and post-chase (r = 0.17, p = 0.0015) periods (Figure 2G). In contrast, plasma cortisol was more strongly correlated with fin cortisol during the pre-chase period, albeit non-significant (r = 0.40, p = 0.1552), as compared to the chase (r = 0.22, 0.05003) and post-chase (r = 0.29, p = 0.0058) period (Figure 2H). Fin and mucus cortisol were most highly correlated in the pre-chase (r = 0.51, p = 0.0643) and chase (r = 0.22, p = 0.1434) period as compared to post-chase (r = 0.09, p = 0.4259) period (Figure 2I).

The dynamics of water cortisol circulating in farm-scale RAS systems has not been previously characterized, posing a practical challenge to implementing on-farm water cortisol monitoring at scale. Hence, we sought to understand how water cortisol may be distributed in the RAS system following an acute stressor. To this end, we quantified cortisol from tank water sampled from 1 to 7 points in the RAS indicated in Figure 1A. In Experiment 1, tank water was only collected from a single sampling point “Tank 1 Furthest from Inlet”, whereas in Experiments 2 and 3, tank water was additionally sampled from 6 other sampling points to study the distribution of cortisol across various locations within the RAS.

Due to the low concentrations of cortisol in water relative to biological samples, and the higher potential for antibody interference in ELISA due to contaminants (e.g., salts and organic material) in tank water, we opted to use HPLC analysis for more precise measurements of water cortisol concentrations. Consistent with reports in existing literature, water cortisol concentrations were in the range of ∼10 to ∼100 ng/L, about 1,000 fold lower than the concentrations observed in plasma and fin samples, and 10-fold lower than mucus samples. In all sampling points within the tank containing the fish (Tank 1), we observed a consistent trend of increased water cortisol starting from the chasing period (Figure 3A-C). For the “Tank 1 Furthest from Inlet” sampling point, there was a significant increase in water cortisol at the Chase (15 min, p = 0.0197) and Post-chase (30 min) time points (p = 0.0367). Water cortisol concentrations were also significantly elevated at the “Inlet of Tank 1) at the Chase (15 min) time point (p = 0.0287), with a similar peak for the “Central pipe” albeit non-significant. The cortisol concentrations from the other RAS compartments did not exhibit distinct variations, although water cortisol concentrations in the “Sump” and “Skimmer” were on average higher during and after the chase (Figure 3D and 3E). The “UV” compartment, on the other hand, did not show any trends (Figure 3F). Interestingly, in Tank 2, which was not holding any fish, a similar trend of increased water cortisol, significant at the Chase (10 min) time point, was observed (Figure 3G). This observation suggests the possibility that cortisol released into the water from one tank may be distributed to other RAS tanks within the system. Detailed cortisol concentration data from individual experiments, as absolute values or normalized to system stocking density (since some fish were removed for sampling), can be found in Supplementary Figures 3-5.

## Discussion

### Handling stress leads to an acute rise in plasma cortisol

In this study, the effects of handling stress on Asian sea bass cortisol concentrations across tissue types were assessed. Figure 4 shows the cortisol concentration ranges found in different tissue types of Asian sea bass and the RAS tank water. Highest concentration of cortisol is observed in plasma, which is very invasive. Less-invasive samples such as fin, mucus, and water have lower cortisol concentration than plasma. Water samples, which are the least invasive, have the lowest concentration range < 300 ng/L.

**Figure 4:**
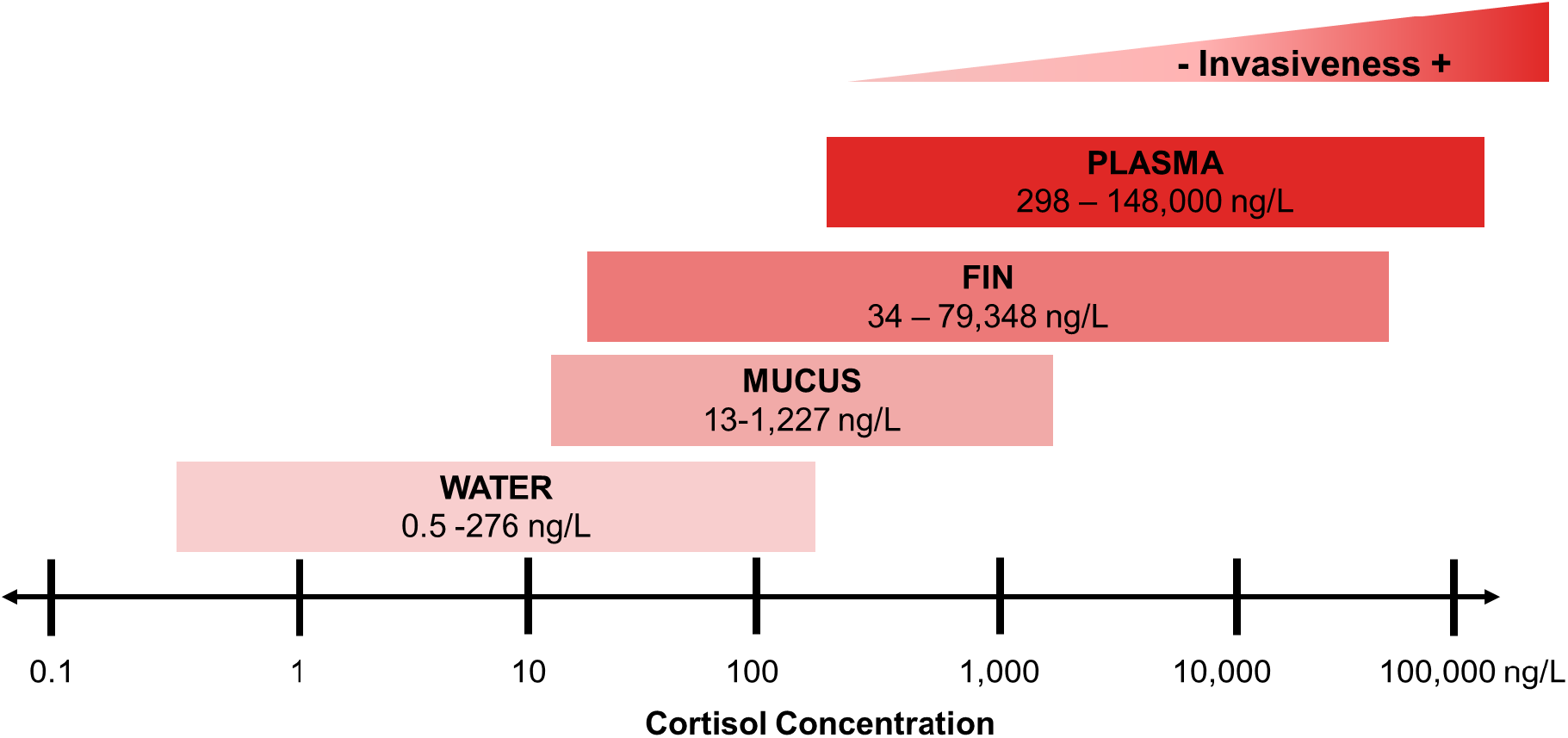
Cortisol concentrations across tissues and in water in Asian sea bass. Cortisol concentrations in this figure are derived from our handling stress experiments. The invasiveness level of each sampling method is indicated by the red color code. Note that the fin and mucus cortisol were normalized to the initial sample weight (100 mg of fin/1 mL of PBS) and mucus was dissolved in 1 mL PBS.

From this acute handling stress study, first, we confirmed that our manipulation indeed triggered a significant increase in plasma cortisol starting from the Chase (10 min) time point, which would be 20 minutes into the chasing period (Figure 2A). There continues to be a significant increase in plasma cortisol up to 1 hour post-chase, and recovery at 24 and 48 hours. We note that some of the chase and post-chase time points did not attain statistical significance, which could be attributed to biological variability across the 3 replicates, influenced by the large number of fish in the tanks and variations in fish age, genetic background, and size (Table 1). Despite these differences, the overall early detection of cortisol is consistent with previous such studies where it has been shown that plasma cortisol typically peaks between 0.5-4 h post stress stimulus in fish (Scott, Pinillos and Ellis, 2001; Fanouraki *et al*., 2008; Scott *et al*., 2008; Mota *et al*., 2017a). For instance, it peaks at 1-2 hr in carp and 2-4 hr in roach post-tag insertion (Lower *et al*., 2005); 2 hr in gilthead seabream post-crowding stress (Guardiola, Cuesta and Esteban, 2016), 0.5 - 1 hr in rainbow trout (Ellis *et al*., 2004) and 3 hr in Atlantic Salmon post air exposure stress (Ellis *et al*., 2007); and 4 hr post-injection with ACTH (adrenocorticotropic hormone) in Threadfin (Bshary *et al*., 2007). Interestingly, a study in European seabass recorded significantly high plasma cortisol already at 0 hr post chasing and air exposure stress, peaking at 1 hr post-stress (Fanouraki *et al*., 2008), which coincides with the plasma cortisol trends in our Asian seabass study.

### Relationship of plasma to mucus and fin cortisol

We also simultaneously sampled mucus and fin cortisol from individual fish, to investigate the viability of these less-invasive sampling methods as reliable indicators of fish stress in aquaculture settings. Our findings revealed that mucus cortisol was significantly increased following the stress-inducing event, becoming significantly elevated at the post-stress 24- and 48-hour time points, even when plasma cortisol had already fully returned to baseline levels. However, this increase followed a slower time course compared to that of plasma cortisol. Similarly as shown in Madaro et al (2022), the rise in cortisol in mucus of Atlantic salmon after netting stress was delayed compared to blood plasma (Madaro *et al*., 2023). In their study, mucus levels remained significantly elevated even at 300 min post-stress, which was the longest time point they assessed. Guardiola et al (2016) also reported a sustained increase in mucus cortisol at 24 and 48-hours post-crowding stress, whereas plasma cortisol only showed a significant increase at 2 hours post-stress (Guardiola, Cuesta and Esteban, 2016).

While the biological basis of these differences is still not well-understood, it is speculated that either the accumulation of plasma cortisol in the mucus, or the reabsorption of cortisol from the water might account for these observed patterns. Despite the differences in cortisol dynamics, we observed a reasonably strong correlation between plasma and mucus cortisol (r = 0.65) during the chasing period. Similarly, one of the few other studies reported the strongest correlation (r = 0.70) between mucus and plasma cortisol in rainbow trout (*Oncorhynchus mykiss*) during the hours following confinement stress, relative to the control or late phase (Carbajal *et al*., 2019). Our data thus suggests that mucus cortisol could be a useful, less invasive way to sample fish stress in Asian Sea bass. In contrast, cortisol extracted from fin did not display any consistent trend or relationship with plasma cortisol. While this does not preclude its utility in estimating fish stress, it may be less robust across the varied conditions in our experiments.

### Water cortisol changes post-stress in the RAS system

We also measured water cortisol within the fish holding tank (Tank 1) across all three acute stress experiments, and observed a significant rise in water cortisol in the Chase (15 min) and post-chase (30 min) periods, which correspond to 40 minutes up to 70 minutes (1 hr 10 mins) from the initiation of the chasing period. In comparison to other previous water cortisol studies, these water cortisol changes were observable in a higher flow system (8000L / hr), again demonstrating the feasibility of water cortisol monitoring in farm-scale settings (Scott, Pinillos and Ellis, 2001; Fanouraki *et al*., 2008; Mota *et al*., 2017a, 2017b). Notably, the rise in water cortisol paralleled the rise in plasma cortisol, except that plasma cortisol continued to be significantly elevated at 1-hr post-chase.

In the latter two of the three experiments, we further measured water cortisol across different parts of the tank and also different RAS compartments, to identify ideal sampling locations for water cortisol for on-farm stress monitoring. The same trend was still observed in other parts of Tank 1, where the peak in water cortisol still occurred at 40 minutes from the commencement of the stress event ( the “Chase (15 min)” time point)) and decreased earlier than plasma cortisol. These results suggest that the dynamics of water cortisol in the RAS system may be influenced by other factors, including flow rate, filtration (i.e. protein skimmer), or binding to the plastic tank walls (since it is hydrophobic). Alternatively, cortisol could potentially be reabsorbed by the fish, possibly into the mucosal layer.

We also observed the presence of water cortisol in other RAS compartments, such as the protein skimmer and sump, indicating the distribution of water cortisol from Tank 1 to other compartments. While water cortisol concentrations in the “Sump” and “Skimmer” were on average higher during and after the chase, the increase was not significant, that could be due to the low number of biological replicates and the varying absolute water cortisol concentrations across the two replicates (Supplementary Figures 3-5). The exception was for the UV compartment, where a change in water cortisol post-chase was not so apparent, either due to UV radiation-induced degradation, or other factors. Interestingly, a rise in cortisol, significant at the Chase (10 min) time point, was observed in Tank 2, which did not contain any fish. This result highlights that due to water recirculation in RAS, other connected tanks may also be exposed to the cortisol released from a stressed tank of fish. It would be interesting to further investigate the effects and implications of water cortisol transmission to fish in connected RAS tanks.

Overall, water cortisol dynamics appeared to be more consistent with plasma cortisol trends than mucus (or fin). We note that we used a more sensitive method (HPLC) to quantify water cortisol than plasma, mucus, or fin cortisol. This was for two reasons, firstly, water cortisol concentrations were significantly lower than in biological samples, and secondly, fish tank water is a more complex matrix than biological samples, containing a multitude of additional salts and organic materials that may interfere with antibody affinity binding in ELISA. More convenient and sensitive methods of water cortisol detection would be valuable for future studies.

### Caveats and limitations of our study

In typical research studies, biological replicates would be conducted in parallel on a single batch of fish, within smaller tank setups. In our study, we aimed to simulate the conditions of a farm-scale setup, using 3,000 L tanks on a 9,000 L RAS system. Due to the scale of our experiments, the limited number of such systems available, and the challenge of acclimatizing and cultivating a large number of fish, we were constrained to conduct one biological replicate at a time, with our experiments spread over a course of 6 months. We also were only able to measure water cortisol from different RAS compartments in 2 of the 3 experiments.

As expected, the cortisol measurements obtained in this study exhibited variability across samples and time points. Several factors contributed to this variability; firstly, the variations in the ages, sizes, genetic background, and stocking densities of the fish tested (Table 1). Secondly, despite maintaining a highly consistent chasing and air exposure stress protocol, the random selection of the 5 fish sampled per time point (per experiment) likely contributed additional variability due to the large numbers of fish present in the tank.

While not ideal in an experimental setting, the upside of such variability is that any signal we have obtained would likely be reproducible even with the normal diversity experienced in on-farm settings. Across all biological replicates, we observed an increase in plasma, mucus, and water cortisol levels, with relatively consistent dynamics. Importantly, our results further argue that surveying water cortisol, which provides an average readout of the stress levels in the system, might be a means of overcoming the variability from sampling individual fish. Further, given that the stocking densities fell within the low-(6 kg/m^3^) to medium-range (20 kg/m^3^), it is possible that more pronounced changes in cortisol might be observable at higher densities.

## Conclusion**s**

Our results corroborate that in a farm-scale RAS setting across varying conditions, water cortisol is a reliable measure of Asian sea bass stress and that different tissue types such as fin and mucus may reflect stress over different dynamical timescales. These results are likely generalizable to other fish species and tank setups, though it is important to note that the time course and concentrations or rates of cortisol release may differ across species. The findings of a significant correlation between mucus cortisol and plasma cortisol, especially during acute stress exposure, highlight the potential utility of mucus cortisol as a non-invasive measure of fish stress. Future experiments could explore the application of these methods to other types of stressors (e.g. ammonia exposure, infections) or more chronic stressors such as high stocking density. Overall, our results suggest that continuous monitoring of water cortisol in aquaculture RAS setups may be a viable approach for detecting fluctuations in fish stress levels, allowing for the detection of anomalies in a timely manner.

## Acknowledgements

This research is supported by the A*STAR Agritech and Aquaculture Horizontal Programme Office (A2HTPO) seed grant (C211018003) awarded to Laura Sutarlie, Xiaodi Su, Caroline Lei Wee and collaborators in Republic Polytechnic and Aquaculture Innovation Center (AIC), and also the Confirma Programme awarded to Caroline Lei Wee by A*STAR. We would like to extend our gratitude to Ng Yi Long, for his valuable help and guidance in fish husbandry. We would also like to acknowledge Tan Kwang Meng and Joyce Koh for their assistance during the experiments. Additionally, we would like to also thank the teams of Republic Polytechnic Final Year Project students: Cung Nei Mawi Lian, Nur Adillah Bte Rahiam, Nur Iffah Dianah Bte Jumadi, Nurul Insyirah Binte Mohamed Arffah, Putri Nur Aisyah Bte Sazali, Yong Sai Kit, Cheryl Lim, Lee Jerry, Sammi Hong Wenxuan and Stahlmann Sarah Elizabeth. We also would like to acknowledge Dr. Lee Chee Wee, Dr. Saravanan Padmanabhan, and Dr. Diana Chan from AIC for discussions.

