## Supplementary Figures and Legends for "Dynamics of endogenous and water cortisol release in Asian Seabass *Lates calcarifer* after acute stress in a farm scale recirculating aquaculture system"

### Supplementary Figure 1

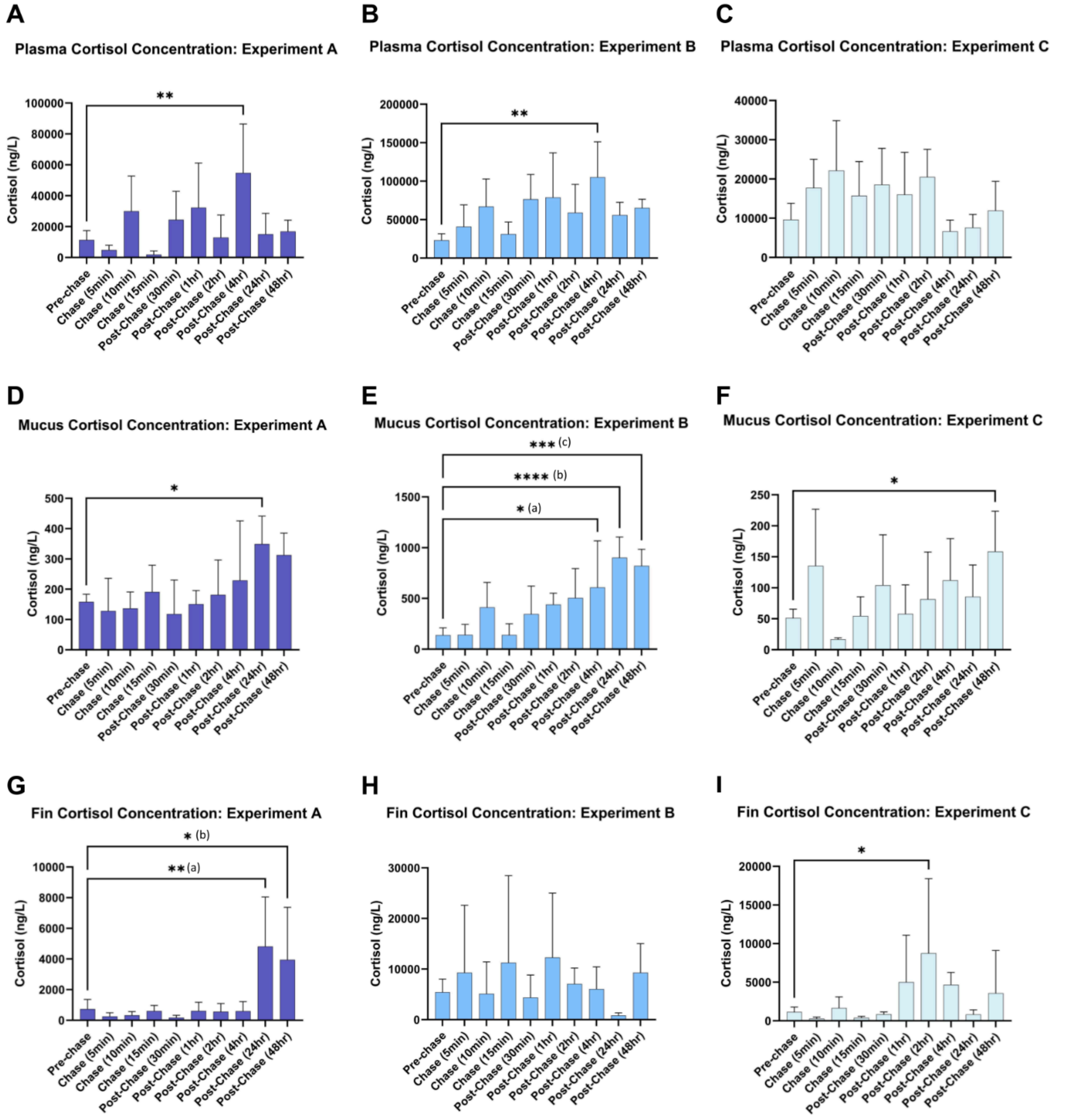


**Supplementary Figure 1: Raw cortisol concentrations in plasma, mucus, and fins across the different sampling points in the RAS system during and after a period of chasing and air exposure stress**

(A) Plasma Cortisol Concentration (ng/L), from experiment A at various sampling points. (N = 5 fish per experiment, 15 fish per sampling point, except for Chase (15min) where one outlier was excluded) p = 0.0039

(B) Plasma Cortisol Concentration (ng/L), from experiment B at various sampling points. (N = 5 fish per experiment, 15 fish per sampling point) p = 0.0023

(C) Plasma Cortisol Concentration (ng/L), from experiment C at various sampling points. (N = 5 fish per experiment, 15 fish per sampling point)

(D) Mucus Cortisol Concentration (ng/L), from experiment A at various sampling points. (N = 5 fish per experiment, 15 fish per sampling point) p = 0.0343

(E) Mucus Cortisol Concentration (ng/L), from experiment B at various sampling points. (N = 5 fish per experiment, 15 fish per sampling point) p = (a) 0.0194, (b) <0.0001, (c) 0.0003,

(F) Mucus Cortisol Concentration (ng/L), from experiment C at various sampling points. (N = 5 fish per experiment, 15 fish per sampling point) p = 0.0433

(G) Fin Cortisol Concentration (ng/L), from experiment A at various sampling points. (N = 5 fish per experiment, 15 fish per sampling point, except for Pre-Chase and Post-Chase (24hr) where one outlier was excluded for each) p = (a) 0.0031, (b) 0.0182

(H) Fin Cortisol Concentration (ng/L), from experiment B at various sampling points. (N = 5 fish per experiment, 15 fish per sampling point)

(I) Fin Cortisol Concentration (ng/L), from experiment C at various sampling points. (N = 5 fish per experiment, 15 fish per sampling point) p = 0.0364

**Supplementary Figure 2**

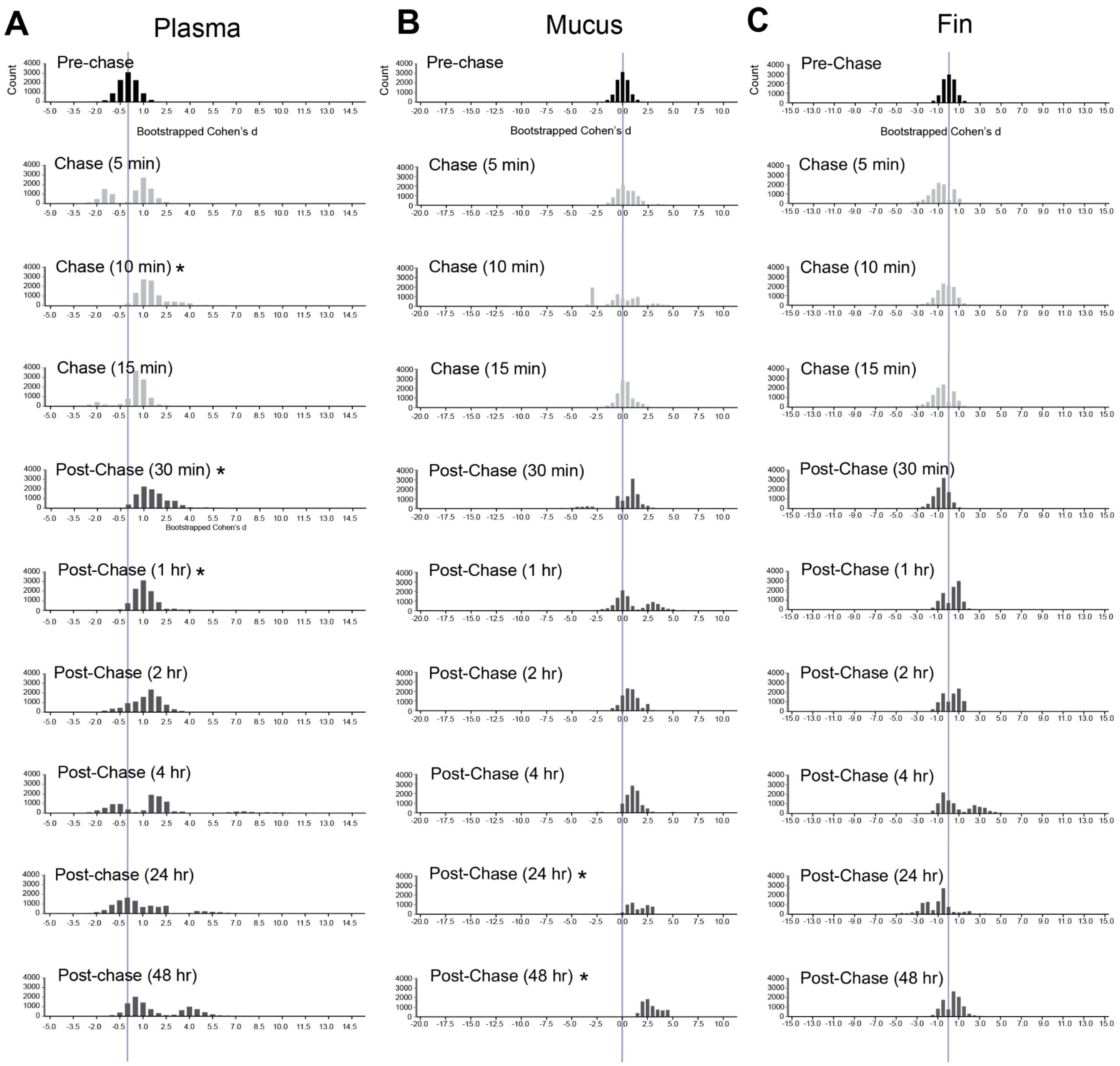


**Supplementary Figure 2: Bootstrapped effect size (Cohen's d) of cortisol concentrations during or post-chase versus pre-chase**

(A) Histograms of bootstrapped Cohen's d (effect sizes) for chase and post-chase plasma cortisol concentrations compared to pre-chase cortisol concentrations. Asterisks indicate time points where concentrations were significantly higher than pre-chase, also reflected by a right-shift in Cohen's d values. Vertical line indicates Cohen's d of zero.

**(B)** Histograms of bootstrapped Cohen's d (effect sizes) for chase and post-chase mucus cortisol concentrations compared to pre-chase cortisol concentrations. Asterisks indicate time points where concentrations were significantly higher than pre-chase, also reflected by a right-shift in Cohen’s d values. Vertical line indicates Cohen’s d of zero.

**(C)** Histograms of bootstrapped Cohen's d (effect sizes) for chase and post-chase fin cortisol concentrations compared to pre-chase cortisol concentrations. Vertical line indicates Cohen’s d of zero.

### Supplementary Figure 3

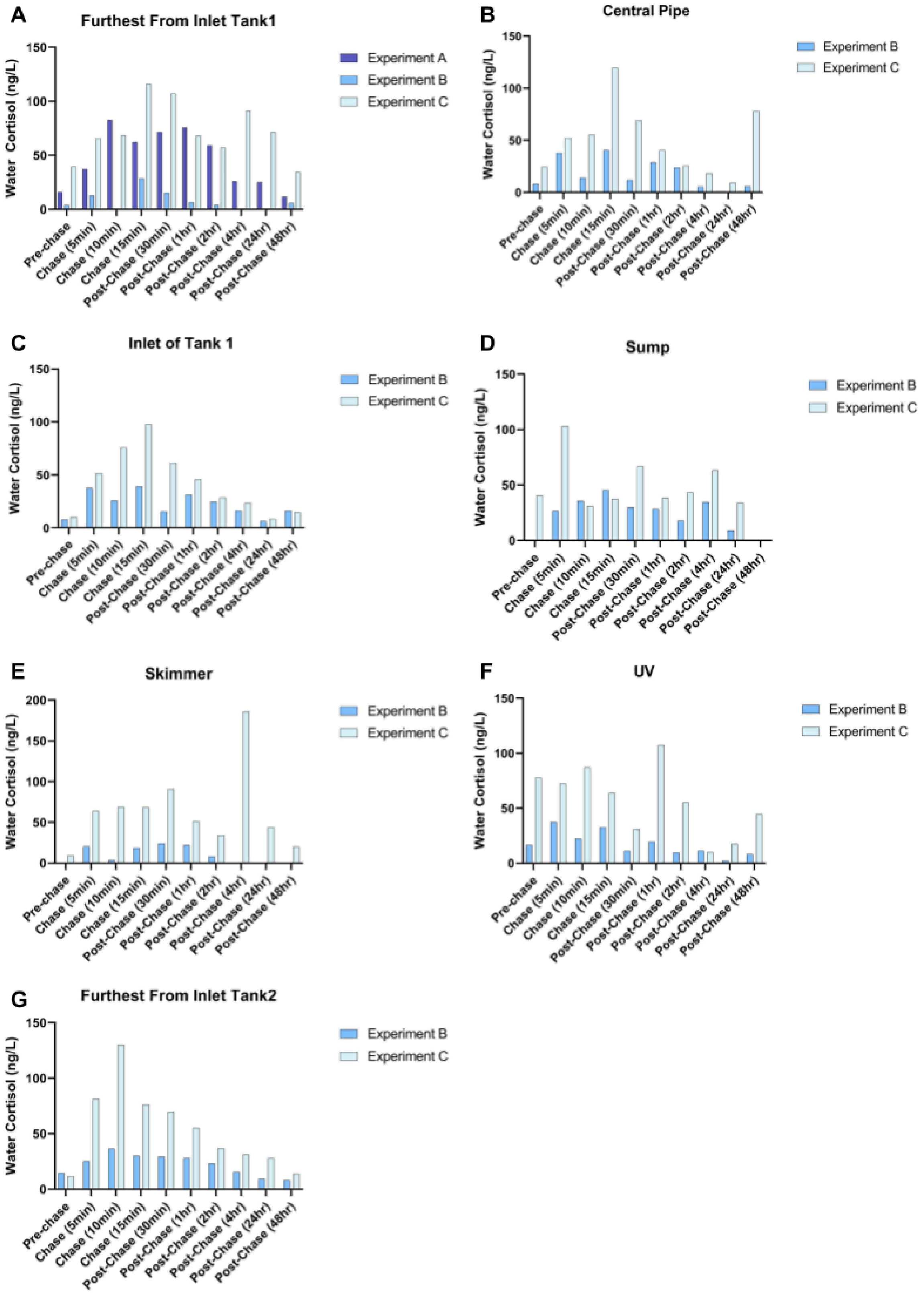

**Supplementary Figure 3: Raw water cortisol concentrations across the different sampling points in the RAS system during and after a period of chasing and air exposure stress**

(A) Water cortisol concentration (ng/L) from Furthest From Inlet Tank 1
(B) Water cortisol concentration (ng/L) from Central Pipe
(C) Water cortisol concentration (ng/L) from Inlet of Tank 1
(D) Water cortisol concentration (ng/L) from Sump Tank
(E) Water cortisol concentration (ng/L) from Skimmer
(F) Water cortisol concentration (ng/L) from UV
(G) Water cortisol concentration (ng/L) from point Furthest From Inlet Tank 2

(N = 1 water sample per collection point)

### Supplementary Figure 4

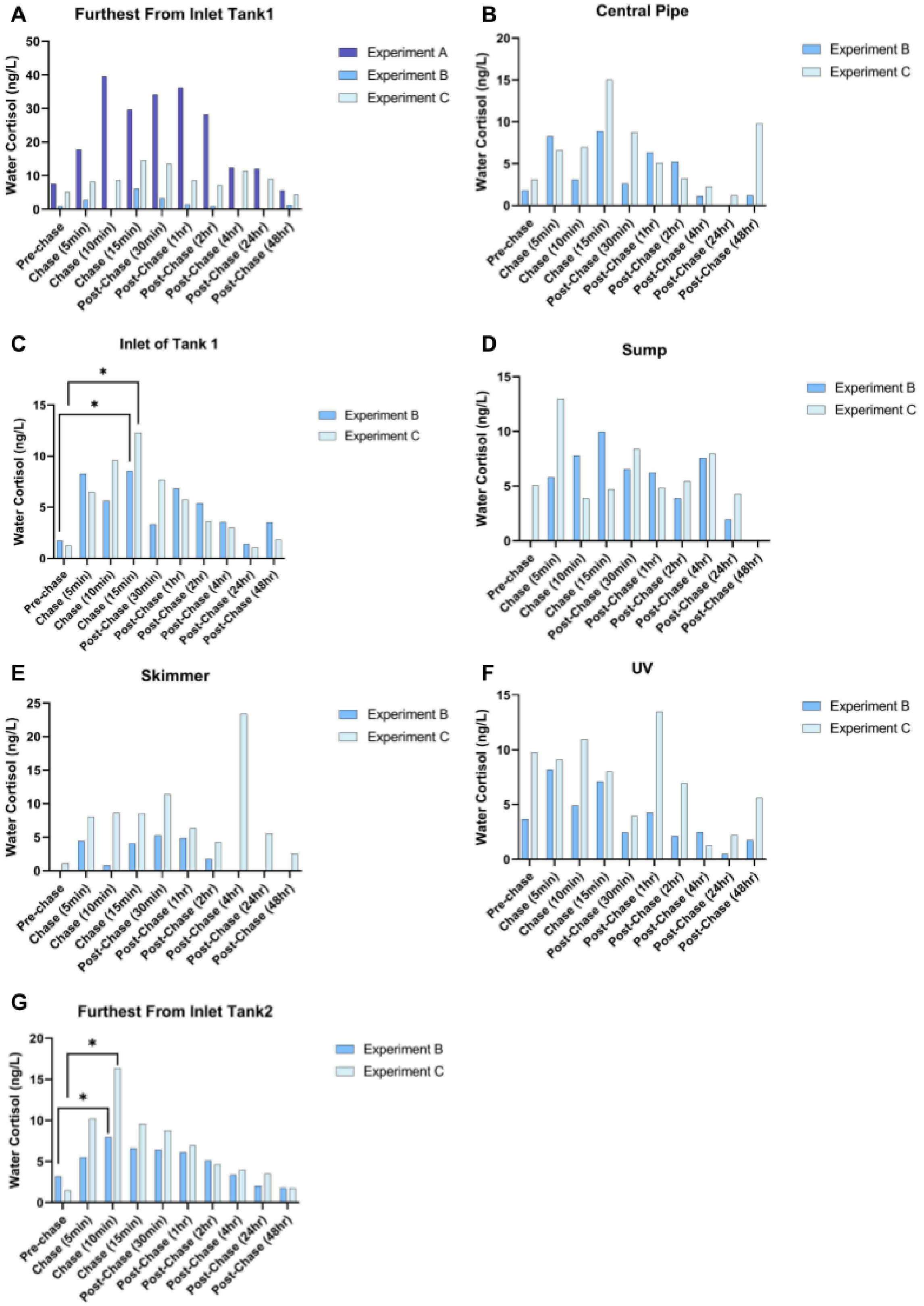

**Supplementary Figure 4: Water cortisol concentrations across the different sampling points in the RAS system during and after a period of chasing and air exposure stress, normalized to system stocking density**

(A) Water cortisol concentration (ng/L) from Furthest From Inlet Tank 1
(B) Water cortisol concentration (ng/L) from Central Pipe
(C) Water cortisol concentration (ng/L) from Inlet of Tank 1 (p = 0.0350 (Pre-chase: Experiment B vs Chase (15min): Experiment B), 0.0350 (Pre-chase: Experiment C vs Chase (15min): Experiment C))
(D) Water cortisol concentration (ng/L) from Sump Tank
(E) Water cortisol concentration (ng/L) from Skimmer
(F) Water cortisol concentration (ng/L) from UV
(G) Water cortisol concentration (ng/L) from point Furthest From Inlet Tank 2 (p = 0.0458 (Pre-chase: Experiment B vs Chase (10min): Experiment B), 0.0458 (Pre-chase: Experiment C vs Chase (10min): Experiment C))

(N = 1 water sample per collection point)

### Supplementary Figure 5

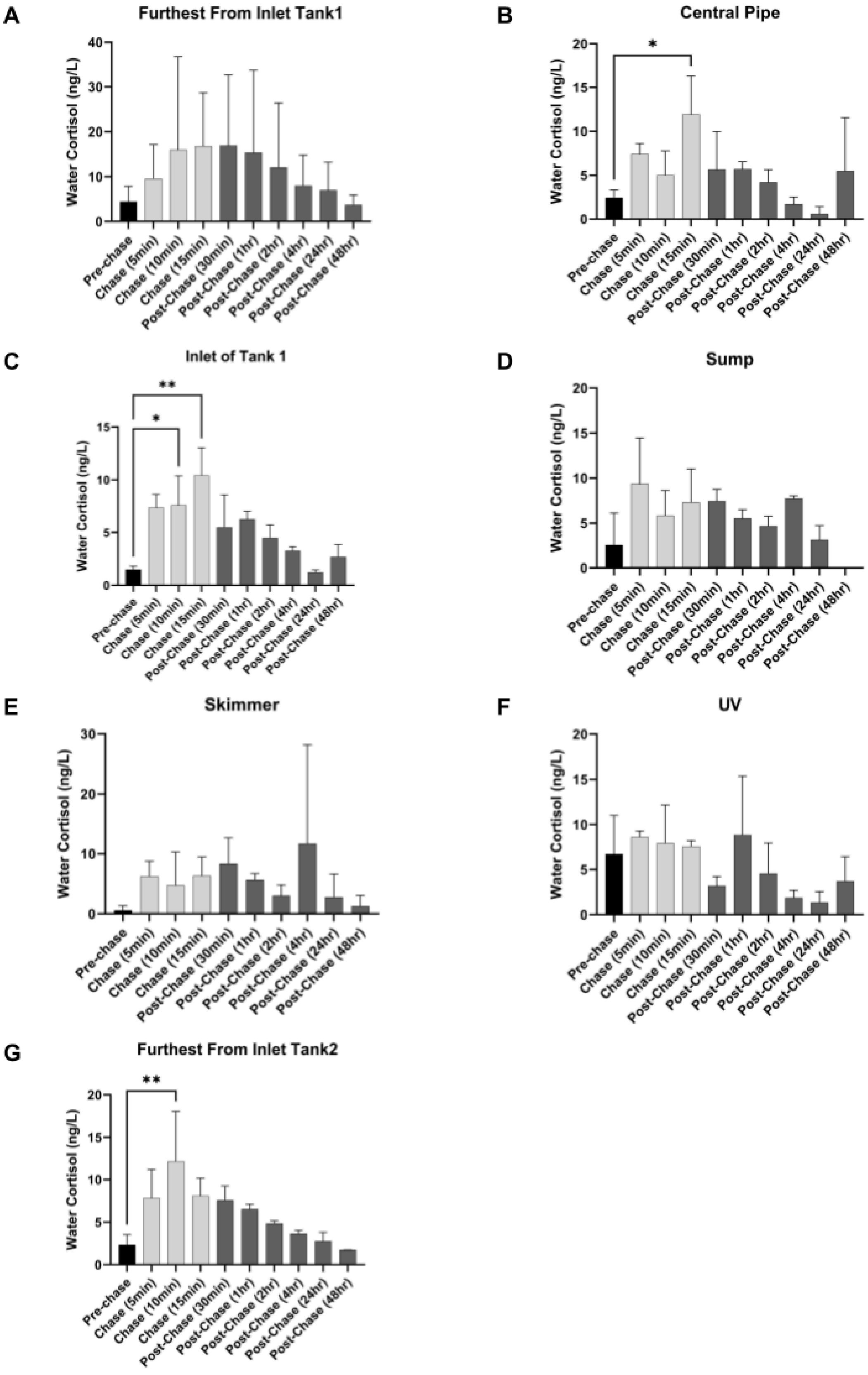

**Supplementary Figure 5: Water cortisol concentrations across the different sampling points in the RAS system during and after a period of chasing and air exposure stress, normalized to system stocking density**

(A) Mean water cortisol concentrations (ng/L) at Furthest From Inlet Tank 1 (N = 3 per time point)
(B) Mean water cortisol concentrations (ng/L) at Central Pipe (N = 2 per time point) p = 0.0302 (Chase (15min))
(C) Mean water cortisol concentrations (ng/L) at Inlet of Tank 1 (N = 2 per time point) p = 0.0432 (Chase (10min)), 0.0046 (Chase (15min))
(D) Mean water cortisol concentrations (ng/L) at Sump Tank (N = 2 per time point)
(E) Mean water cortisol concentrations (ng/L) at Skimmer (N = 2 per time point)
(F) Mean water cortisol concentrations (ng/L) at UV (N = 2 per time point)
(G) Mean water cortisol concentrations (ng/L) at Furthest From Inlet Tank 2 (N = 2 per time point) p = 0.0063 (Chase (10min))
