## Supplementary Table 1 for "Dynamics of endogenous and water cortisol release in Asian Seabass *Lates calcarifer* after acute stress in a farm scale recirculating aquaculture system"

| **Experiment** | **Stocking Density (Tank 1)** | | | | |
| --- | --- | --- | --- | --- | --- |
|  | Pre-chase | Post-chase  (30 min) | Post-chase  (4 hr) | Post-chase  (24 hr) | Post-chase (48 hr) |
| 1 | 6.3 | 5.1 | 4.65 | 4.4 | 4.2 |
| 2 | 13.7 | 12.9 | 12.4 | 12.3 | 12.1 |
| 3 | 23.9 | 22.5 | 21.7 | 21.4 | 21.1 |
| **Experiment** | **Stocking Density (System)** | | | | |
|  | Pre-chase | Post-chase  (30 min) | Post-chase  (4 hr) | Post-chase  (24 hr) | Post-chase  (48 hr) |
| 1 | 2.1 | 2.1 | 2.1 | 2.1 | 2.1 |
| 2 | 4.6 | 4.3 | 4.15 | 4.6 | 4.6 |
| 3 | 8.0 | 7.4 | 7.2 | 7.1 | 7.0 |

**Supplementary Table 1:** Stocking density changes in the system or tank across different experimental time points. While there was an inevitable reduction in tank stocking density due to fish sampling, there were smaller changes to system stocking density, hence water cortisol concentrations should be minimally affected (water cortisol normalised to system stocking density can be found in Supplementary figures 3-4)
